# The effects of Arabian jasmine on zebrafish behavior depends on strain, sex, and personality

**DOI:** 10.1101/2025.05.16.654482

**Authors:** Tripatchara Atiratana, Aliyah R. Goldson, Siritron Samosorn, Neha Rajput, Nalena Praphairaksit, Justin W. Kenney

## Abstract

*Jasminum sambac* (L.) Aiton, commonly known as Arabian jasmine, is widely used in Thai traditional medicine for mental health ailments. While most studies in humans and animals find that Arabian jasmine reduces stress and anxiety, there are a handful of reports that it can oppose relaxation by increasing autonomic arousal. Using adult zebrafish, we sought to determine whether factors like strain, sex, and personality might contribute to the variable effects of *J. sambac* on anxiety-related behavior. The flowers of *J. sambac* were extracted by ultrasonic-assisted extraction with optimal air pressure. Headspace solid-phase microextraction with gas chromatography-mass spectrometry (HS-SPME-GC-MS) identified the main components in the Arabian jasmine flower extract, including linalool (an anxiolytic compound) and benzaldehyde. We fed three strains of zebrafish (AB, TL, and WIK) a gelatin pellet containing different concentrations of *J. sambac* (5-20 mg kg^-1^) and assessed 3-dimensional swim behavior in the novel tank and mirror biting tests. We found that in female AB fish, *J. sambac* resulted in a decrease in bottom distance during the novel tank test, consistent with an anxiogenic effect; there was no effect in WIK or TL fish. We also found that behavioral/personality type influenced the effects of *J. sambac* where shy AB females increased their percent explored, consistent with an anxiolytic effect. Thus, we find that sex, genetics, and personality interact to influence the anxiety-related effects of Arabian jasmine suggesting that these factors may contribute to the opposing effects of Arabian jasmine reported in the literature.

## Introduction

*Jasminum sambac* (L.) Aiton (common name “Arabian jasmine”, Thai name “Ma-li-la”) is a plant that is commonly used in the traditional medicine of many countries. For example, in Thailand, this plant species is contained in the Ya Hom Thep Pa Chit preparation, which is made of 50% Arabian jasmine flowers and consumed orally as an infusion as an anxiolytic [1]. Indeed, Arabian jasmine has been found to result in an increase in self-reported levels of relaxation and reduced levels of stress hormones [2–4], consistent with an anxiolytic effect. However, there are also reports that Arabian jasmine can act to stimulate arousal [5,6], which would oppose relaxation. Thus, it is unclear how Arabian jasmine can have these two opposing properties. It may be that biological factors of participants, like sex, background genetics, and personality, may influence an individual’s responses to Arabian jasmine.

Arabian jasmine is rich in phytochemicals, with benzaldehyde and linalool being particularly prominent [7,8]. Linalool, which has a floral or spicy-wood scent, has been found to reduce aggression and anxiety-related behaviors [3,9,10]. Benzaldehyde is characterized by an almond-like odor and, and although its behavioral effects have not been as well studied, there is at least one report of it increasing aggression [11]. These compounds, alongside several others, create a complex chemical composition that is affected by specific cultivation practices that could contribute conflicting findings regarding the behavioral and autonomic effects of Arabian jasmine.

Individuals can differ in their behavioral and physiological responses to compounds due to variation in factors such as sex, genetics, and even personality traits. This differential sensitivity provides a potential explanation for paradoxical drug effects (i.e., outcomes opposite to those expected) observed with some drugs. For example, sex can influence how animals respond to drugs like alcohol [12,13]. Genetic variation has long been known to modulate the effects of psychoactive drugs in both model organisms and humans, giving rise to the field of pharmacogenomics [14]. Finally, personality-like traits have also been found to influence the behavioral effects of drugs [15,16]. Here, we hypothesized that the anxiety-related effects of Arabian jasmine will be affected by a mix of sex, genetics, and personality. To test this hypothesis, we examined the effects of Arabian jasmine in zebrafish on anxiety-related behaviors during exploration of a novel environment.

Zebrafish (*Danio rerio* (Hamilton,1822)) are a tropical freshwater fish native to south Asia [17] that has grown in popularity as a model organism in behavioral research over the past several decades [18–20]. This popularity is due to their small size and affordability coupled with high genetic similarity to humans [21]. Zebrafish also possess all the main neurotransmitter systems as mammals [22] with overlapping molecular mechanisms [23], and stress hormones [24]. Finally, adult zebrafish exhibit a wide variety of behaviors [25] making them a viable animal model for behavioral and pharmacological studies. With respect to sex, genetics, and personality, recent studies have found that, like mammals, zebrafish behavior is influenced by these factors [26,27], and that they display consistent individual differences in behavior (i.e., personality) [27,28]. Here, we find that female zebrafish of the AB (but not WIK or TL) strain increase their anxiety-like behavior in response to Arabian jasmine. We also find that zebrafish personality type (boldness and activity) affect the behavioral response to Arabian jasmine. Our finding indicates that sex, genetic variation and personality may play an important role in the paradoxical effects of Arabian jasmine on anxiety-related behavior.

## Results

### Chemical profile of Arabian jasmine

Arabian jasmine (Fig. 1A) was extracted via ultrasonic-assisted extraction with air pressure (Fig. 1B), using water as the solvent. Its composition was determined using Headspace Solid-Phase Microextraction Coupled with Gas Chromatography–Mass Spectrometry (HS-SPME-GC-MS) (Fig. 1C and Table S1). Among the 40 molecules identified, the top two comprised over half the sample (dimethyl sulfide: 30.2% and linalool: 28.4%). Four of the identified compounds have previously been reported to have anxiolytic properties: 3-hexen-1-ol (0.5%) [29], linalool [3,9,10,30–32], linalool oxide (0.09%) [33], phenylethyl alcohol (1.46%) [34]. Linalool is notable because of its high concentration and its reported behavioral and psychoactive effects [3,9,10,30–32]. In addition, we also identified small levels of other compounds, like benzaldehyde (0.68%), that has been reported to have potentially anxiogenic effects [11].

**Fig. 1.**
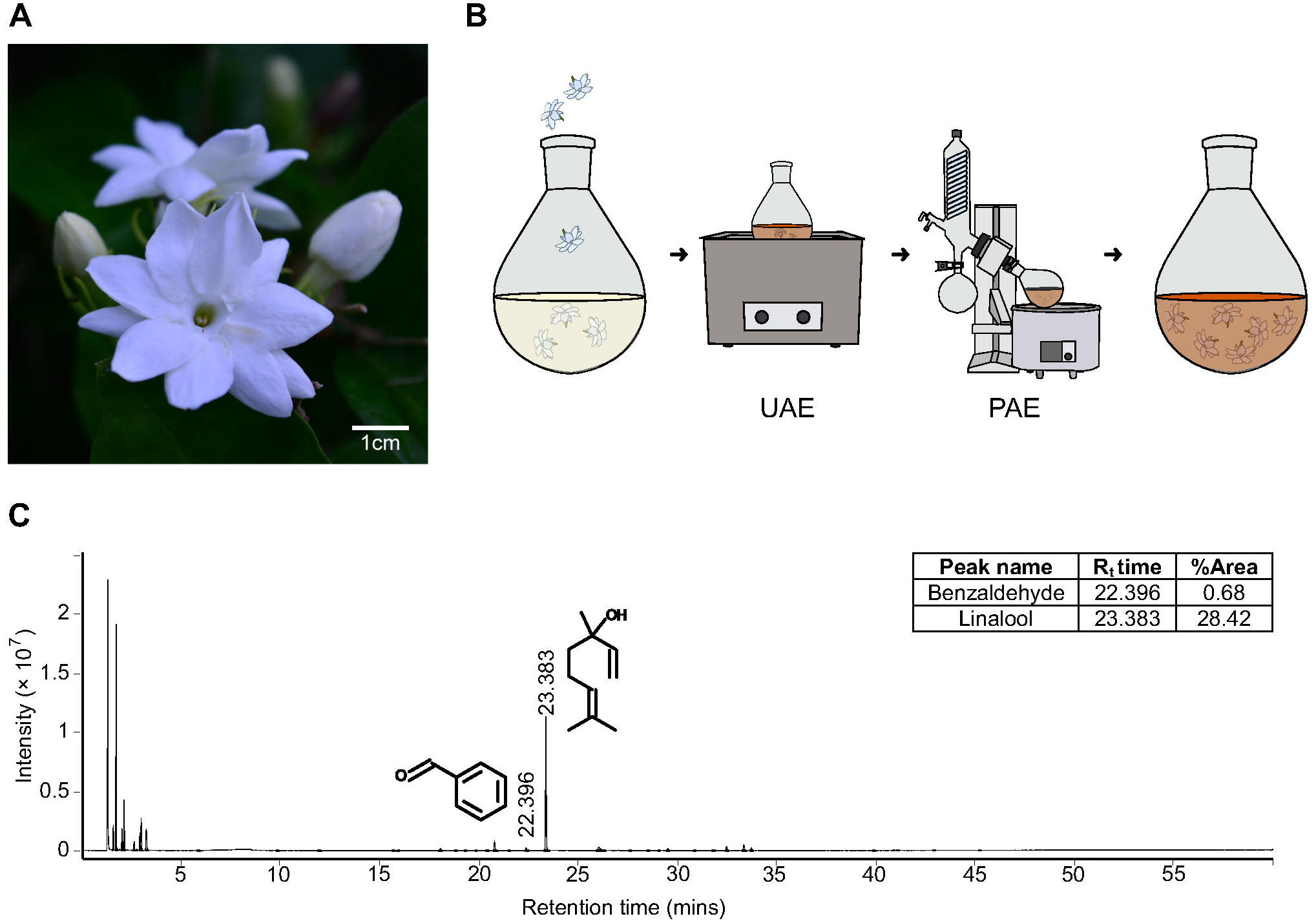
The chemical studies of Arabian jasmine flower. (A) Example image of Arabian jasmine flowers used for this study. (B) Experimental design for extracting Arabian jasmine flowers as done using a traditional approach. Flowers were extracted in water by ultrasonic-assisted extraction (UAE) followed by pressure-assisted extraction (PAE). (C) GC chromatogram with MS spectra of Arabian jasmine showing linalool and benzaldehyde. Retention time (R_t_ time) is the amount of time a compound spends in the GC column before being detected by MS. %Area indicates the relative amount of each compound based on the size of its peak in the GC chromatogram.

### Sex and genetics influence the effects of Arabian jasmine on exploration of a novel tank

To determine if sex and genetics influence the behavioral effects of Arabian jasmine, we did a dose response (0, 5, 10, 20 mg kg^-1^) in two strains of fish (AB and WIK) and both sexes in novel tank test (NTT) and mirror biting test (MBT). Behavior in the NTT was used to identify changes in anxiety-like behaviors (e.g., bottom distance) and locomotor activity, and the MBT was used to identify changes in aggressive behaviors. We assessed significance using 4 × 2 (dose × sex) ANOVAs followed by Dunnett’s multiple comparison tests with the vehicle as the control group.

For the NTT, we assessed the effects of Arabian jasmine on bottom distance, center distance, percentage of tank explored, and distance travelled (Fig. 2A and Table S2). For bottom distance in AB zebrafish (Fig. 2A), we found a trend towards a small effect of dose (*P*=0.085, η^2^=0.06), a trend towards a small interaction (*P=*0.072, η^2^=0.06), and no effect of sex (*P*=0.50). Post-hoc tests indicated large effects in the 10 mg kg^-1^ (*P=*0.010, *d*=1.22) and 20 mg kg^-1^ (*P=*0.038, *d*=1.18) female treatment groups where Arabian jasmine treated fish swam closer to the bottom of the tank than the vehicle group. For the bottom distance in WIK zebrafish (Fig. 2A), we found no effect of dose (*P=*0.11), sex (*P=*0.80), or an interaction (*P=*0.59). For the center distance in AB zebrafish (Fig. 2A), we found a trend towards a small effect of sex (*P=*0.098, ^2^=0.02), but no effect of dose (*P=*0.88), and no interaction (*P=*0.47). Post-hoc tests did not show a significant difference from the control group. For the center distance in WIK zebrafish (Fig. 2A), we found no effect of dose (*P=*0.87), sex (*P=*0.22), or an interaction (*P=*0.16). For the percentage of exploration in AB zebrafish (Fig. 2A), we found a trend towards a small effect of dose (*P* =0.064, η^2^=0.06), but no effect of sex (*P=*0.29), or an interaction (*P=*0.51). Post-hoc tests indicated that female fish eating 10 mg kg^-1^ had a trend towards a large effect of less exploration compared to the vehicle group (*P=*0.063, *d*=0.90). For the percentage of exploration in WIK zebrafish (Fig. 2A), we found no effect of dose (*P=*0.60), sex (*P=*0.92), or an interaction (*P=*0.86). Finally, for the distance travelled, in AB zebrafish (Fig. 2A), we found a large effect of sex (*P<*0.001, η^2^=0.20), where males swam further than females, but no effect of dose (*P=*0.91), or an interaction (*P=*0.74). In WIK zebrafish (Fig. 2A), we also found a large effect of sex (*P<*0.001, η^2^=0.16), with males travelling further than females, but no effect of dose (*P=*0.28), or an interaction (*P=*0.42).

**Fig. 2.**
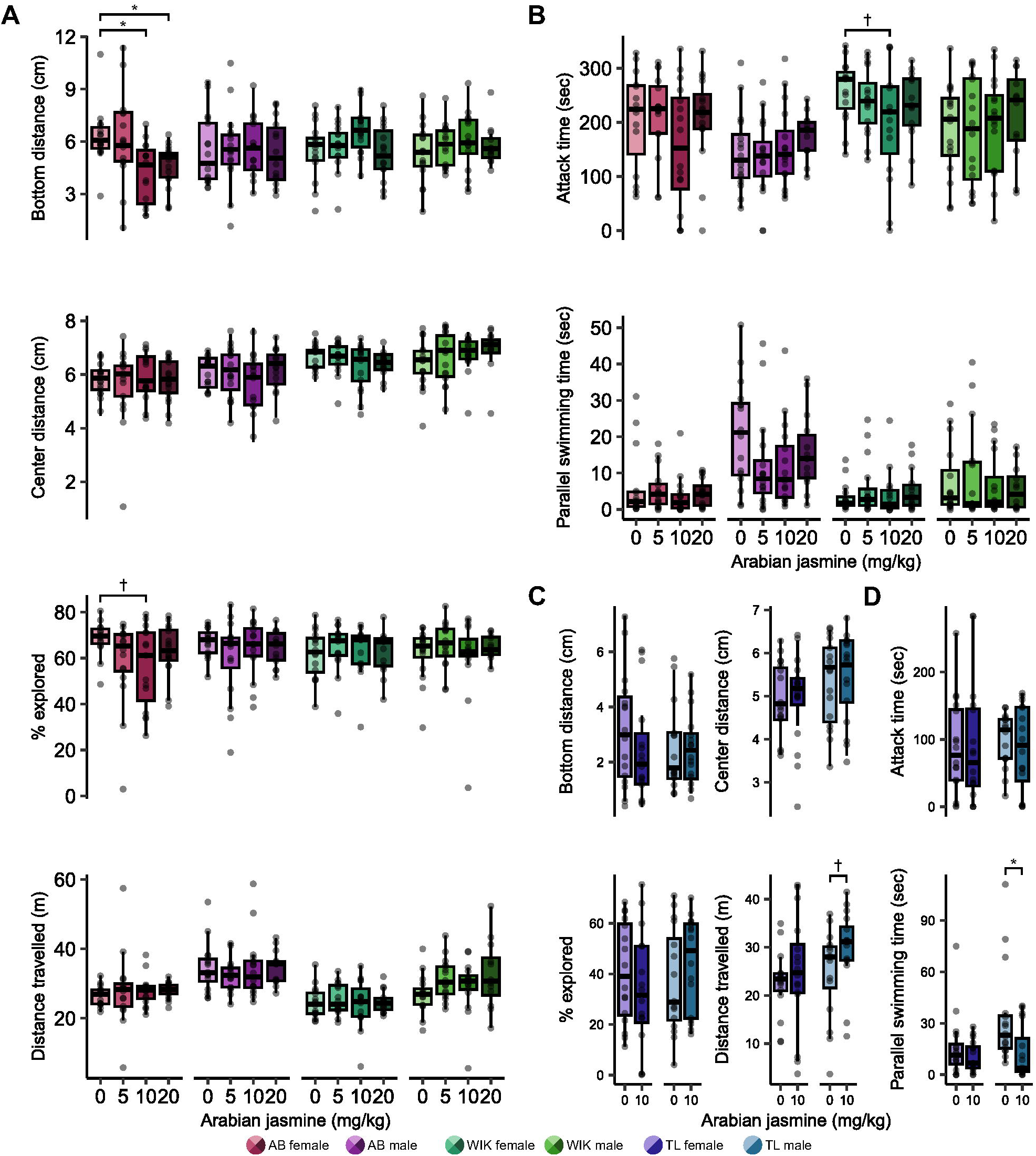
Influence of sex and strain on the behavioral effects of Arabian jasmine. The effect of Arabian jasmine on (A) exploratory behaviors (bottom distance, center distance, percent of exploration, and distance travelled) and (B) social behaviors (attack time and parallel swim time) in AB and WIK zebrafish based on dose (0, 5, 10, 20 mg kg^-1^) and sex. The effect of Arabian jasmine on (C) exploratory behaviors and (D) social behaviors in TL zebrafish given either vehicle or 10 mg kg^-1^ Arabian jasmine. Boxplots indicate median (center line), interquartile range (box ends), and hinge±1.5 times the interquartile range (whiskers). *P<0.05, †P<0.10 from Dunnett’s post-hoc in comparison to the 0 mg kg^-1^ group (Fig. 2A-B). *P<0.05, †P<0.10 from pairwise *t*-test with FDR correction within group (Fig. 2C-D). AB female: *n*=15 vehicle (0 mg kg^-1^ Arabian jasmine); *n*=16 5 mg kg^-1^ Arabian jasmine; *n*=16 10 mg kg^-1^ Arabian jasmine; *n*=16 20 mg kg^-1^ Arabian jasmine. AB male: *n*=16 vehicle; *n*=16 5 mg kg^-1^ Arabian jasmine; *n*=16 10 mg kg^-1^ Arabian jasmine; *n*=14 20 mg kg^-1^ Arabian jasmine. WIK female: *n*=16 vehicle; *n*=16 5 mg kg^-1^ Arabian jasmine; *n*=15 10 mg kg^-1^ Arabian jasmine; *n*=15 20 mg kg^-1^ Arabian jasmine. WIK male: *n*=16 vehicle; *n*=16 5 mg kg^-1^ Arabian jasmine; *n*=16 10 mg kg^-1^ Arabian jasmine; *n*=12 20 mg kg^-1^ Arabian jasmine. TL female: *n*=16 vehicle; *n*=16 10 mg kg^-1^ Arabian jasmine. TL male: *n*=16 vehicle; *n*=16 10 mg kg^-1^ Arabian jasmine.

During the MBT we assessed both attack and parallel swim time (Fig. 2B and Table S3). For the attack time in AB zebrafish (Fig. 2B), we found a medium-sized effect of sex (*P=*0.004, η^2^=0.07), where females attacked more than males, but no effect of dose (*P=*0.36), or an interaction (*P=*0.22). For attack time in WIK zebrafish (Fig. 2B), we again found a small effect of sex (*P=*0.014, ^2^=0.05), but no effect of dose (*P=*0.59), or an interaction (*P=*0.37). Finally, for the parallel swimming time in AB zebrafish (Fig. 2B), we found a large effect of sex (*P<*0.001, η^2^=0.22), where males engaged in more parallel swimming than females, and a trend towards a small effect of dose (*P=*0.094, ^2^=0.05), but no interaction (*P=*0.44). However, post-hoc tests did not find any significant differences from the vehicle control in either males or females. For parallel swimming time in WIK zebrafish (Fig. 2B), we found a trend towards a small effect of sex (*P=*0.066, η^2^=0.03), but no effect of dose (*P=*0.63), or an interaction (*P=*0.83).

From the results of the dose-response experiment in NTT and MBT, we found that the 10 mg kg^-1^ treatment group had an anxiogenic effect in AB zebrafish but no effect in WIK zebrafish, thus, we hypothesized that this dose could affect the anxiety-related behavior in zebrafish based on genetic variation. Thus, we examined the effect of 10 mg kg^-1^ Arabian jasmine on zebrafish of the TL strain as this strain has been found to have higher baseline levels of anxiety-like behavior than other strains [27]. For bottom distance in TL zebrafish, we found no effect of Arabian jasmine (*P=*0.49), sex (*P*=0.43), and no interaction (*P*=0.38). For the center distance of TL zebrafish, we found a trend towards a small effect of sex (*P*=0.089, ^2^=0.05), but no effect of Arabian jasmine (*P*=0.79), or an interaction (*P*=0.88). For the percentage of exploration in TL zebrafish, we found no effect of Arabian jasmine (*P*=0.97), sex (*P*=0.52), and there was no interaction (*P*=0.17). For the distance travelled of TL zebrafish, we found no effect of Arabian jasmine (*P*=0.12), sex (*P*=0.12), and no interaction (*P*=0.52). However, post-hoc tests found that the 10 mg kg^-1^ male treatment group (*P*=0.10, *d*=0.59) had a trend towards a medium-sized effect of higher activity than the vehicle group.

For the attacking time of TL zebrafish, we found no effect of Arabian jasmine (*P=*0.77), sex (*P=*0.89), and no interaction (*P=*0.68). For the parallel swimming time of TL zebrafish, we found a medium-sized effect of Arabian jasmine (*P=*0.009, η^2^=0.11), a trend towards a medium-sized effect of sex (*P=*0.051, ^2^=0.06), but no interaction (*P=*0.12). Post-hoc tests found that the 10 mg kg^-1^ male treatment group (*P=*0.017, *d*=0.89) had less parallel swimming time than the vehicle treated group.

### The effect of Arabian jasmine on anxiety-related behaviors is influenced by zebrafish personality

From the genetic variation experiment, we found that TL and WIK zebrafish had no response to Arabian jasmine in the NTT, whereas AB zebrafish had an anxiogenic response at 10 mg kg^-1^. We noticed a wide variation in behavioral response to Arabian jasmine in the AB fish and hypothesized that perhaps the personality of the animals may affect how they respond to Arabian jasmine. To test this, we examined two different personality-related behaviors: boldness as measured by a combination of bottom distance and percent of tank explored [15,27], and activity via distance travelled [28]. To determine personality type, we first performed the NTT in the absence of Arabian jasmine on day 1, then, on day 2, individuals were given 10 mg kg^-1^ Arabian jasmine to determine their response in the NTT and MBT. We assessed significance using 2 × 2 (treatment × boldness/activity level) ANOVAs within each sex followed by FDR-corrected pair-wise *t*-tests to compare groups. One-sample *t-*tests were used to determine if the behavior changed from day 1 to day 2. The distribution of the boldness index in each sex is presented in Fig. 3A and Table S4, and the distribution of the activity index in each sex is presented in Fig. 3B. We found no correlation between activity and boldness, suggesting these measures are capturing distinct aspects of zebrafish behavior (Fig. 3C).

**Fig. 3.**
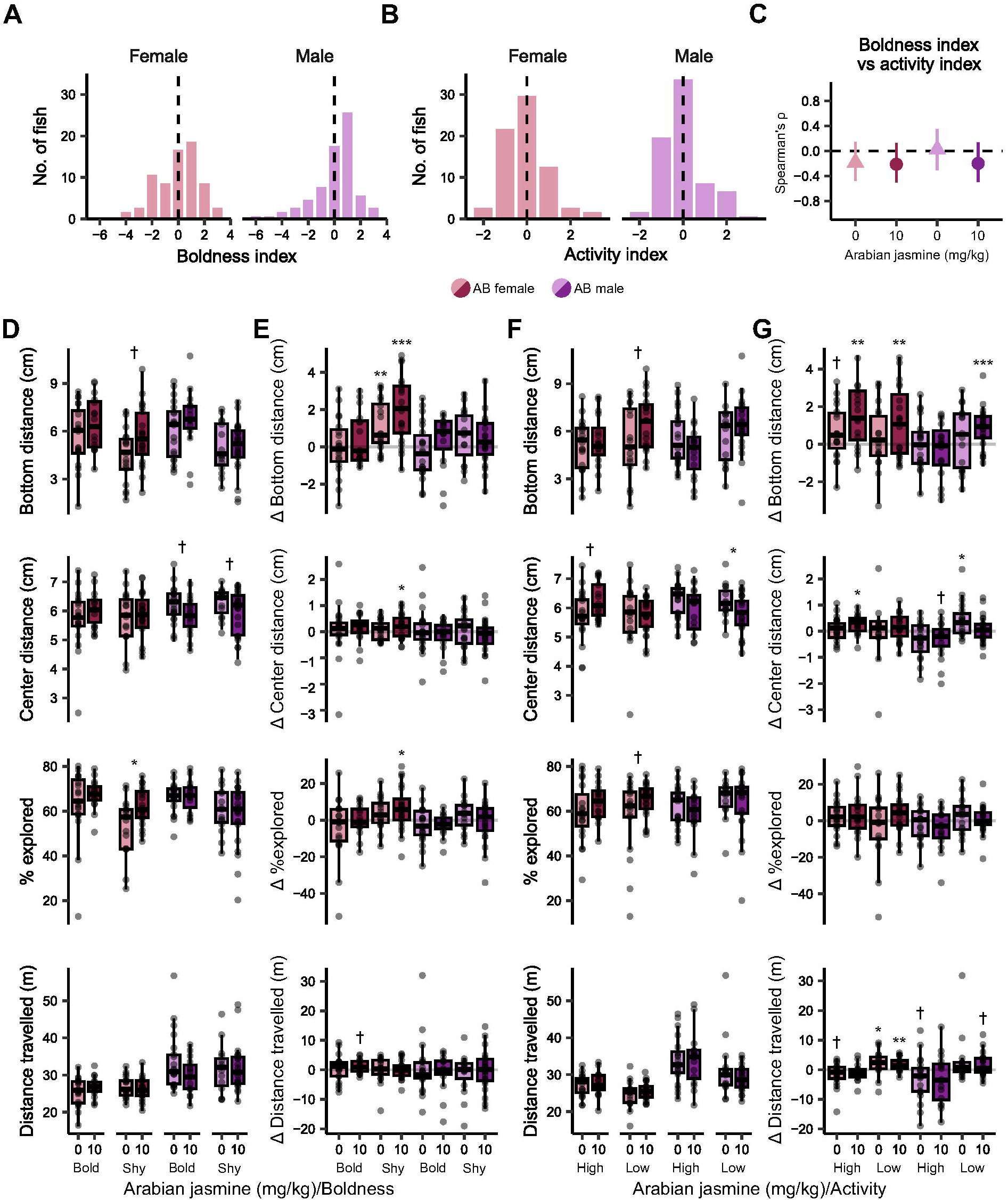
Influence of personality of the behavioral effects of Arabian jasmine. (A) Histogram of boldness index, and (B) activity index on day 1 of exposure to the novel tank. The dashed line is the median. (C) Spearman’s rank correlation coefficient (ρ) between boldness index and activity index; error bars are 95% confidence intervals. Exploratory behaviors on day 2 based on (D) boldness and (F) activity. Fish were given either vehicle or 10 mg kg^-1^ Arabian jasmine. Change in behavior between the first and second days of exploratory behaviors based on (E) boldness and (G) activity and Arabian jasmine administration. Boxplots indicate median (center line), interquartile range (box ends), and hinge±1.5 times the interquartile range (whiskers). *P<0.05, †P<0.10 from pairwise t-test with FDR correction within group. ***P<0.001, **P<0.01, *P<0.05, †P<0.10 compared to zero using one-sample t-tests with FDR corrections. Bold female: *n*=19 vehicle; *n*=16 Arabian jasmine. Shy female: *n*=17 vehicle; *n*=20 Arabian jasmine. Bold male: *n*=21 vehicle; *n*=15 Arabian jasmine. Shy male: *n*=15 vehicle; *n*=22 Arabian jasmine. High-active female: *n*=19 vehicle; *n*=16 Arabian jasmine. Low-active female: *n*=17 vehicle; *n*=20 Arabian jasmine. High-active male: *n*=20 vehicle; *n*=16 Arabian jasmine. Low-active male: *n*=16 vehicle; *n*=21 Arabian jasmine.

First, we determined if boldness influences the behavioral response of zebrafish to Arabian jasmine (Fig. 3D-E; Fig. S1A, Tables S5 and S6). For the bottom distance of AB female zebrafish, we found a medium effect of Arabian jasmine (*P=*0.045, η^2^=0.06), and boldness (*P=*0.030, ^2^=0.07), but no interaction (*P=*0.60). Post-hoc tests found that the Arabian jasmine in shy females fish had a trend towards a small anxiolytic effect where they increased their bottom distance (*P=*0.056, *d*=0.65). For AB male zebrafish, we found a large effect of boldness on bottom distance (P=0.0011, η^2^=0.14), but no effect of Arabian jasmine (*P*=0.86), or an interaction (*P*=0.47). For the change in bottom distance over time, a one-sample *t*-test found that only shy female zebrafish, irrespective of drug treatment, increased distance from bottom (*P*=0.005, *d*=0.69 for vehicle treated, and *P*<0.001, *d*=1.24 for Arabian jasmine treated).

For center distance, in female AB zebrafish, we found no effect of Arabian jasmine (*P*=0.36), boldness (*P*=0.80), and no interaction (*P*=0.56). In male AB zebrafish, we found a medium effect of Arabian jasmine (*P*=0.017, η^2^=0.08), but no effect of boldness (*P*=0.61), or an interaction (*P*=0.98). Post-hoc tests found a trend toward medium size effects in Arabian jasmine treated bold males (*P*=0.070, *d*=0.63) and shy males (*P*=0.100, *d*=0.56), both swimming closer to the center of the tank. For the change in center distance over time, a one-sample t-test found that the Arabian jasmine shy female treatment group swam closer to the border of the tank on the second day (*P*=0.049, *d*=0.30).

For the percentage of exploration of female AB zebrafish, we found a medium sized effect of Arabian jasmine (*P*=0.030, η^2^=0.07), and boldness (*P*=0.007, η^2^=0.10), but no interaction (*P*=0.40). Post-hoc tests found that Arabian jasmine caused shy female fish to explore the tank more than the vehicle group (*P*=0.016, *d*=0.84). For AB male zebrafish, we found a medium-sized effect of boldness (*P*=0.007, η^2^=0.10), but no effect of Arabian jasmine (*P*=0.38), or an interaction (*P*=0.59). For the change in percentage of exploration over time, the Arabian jasmine treated shy female group explored the tank more than on day 1 (*P*=0.029, *d*=0.80).

For distance travelled, in AB female zebrafish we found no effect of Arabian jasmine (*P*=0.66), boldness (*P*=0.56), and no interaction (*P*=0.17). For distance travelled in males, we also found no effect of Arabian jasmine (*P*=0.31), boldness (*P*=0.91) and no interaction (*P*=0.20). For the change in distance travelled over time, we found that Arabian jasmine in bold females had a trend towards more activity than on day 1 (*P*=0.062, *d*=0.45).

Next, we determined if activity levels influence the behavioral effects of Arabian jasmine in the NTT and MBT (Fig. 3F-G, S1B and Table S6). There were no significant effects in the MBT. In the NTT, for bottom distance in female zebrafish, we found a medium sized effect of Arabian jasmine (*P*=0.046, η^2^=0.06), but no effect of activity (*P*=0.10), and no interaction (*P*=0.25). Post-hoc tests found that the low-activity female treatment group had a trend towards a medium-sized effect of bottom distance being higher than the vehicle group (*P*=0.054, *d*=0.66). For males, we found a medium-sized effect of activity (*P*=0.025, η^2^=0.07), a trend towards a small interaction (*P*=0.076, η^2^=0.05), but no effect of treatment (*P*=0.86). However, post-hoc tests found no differences between groups. For the change in bottom distance over time, we found an effect of Arabian jasmine in females in both the high (*P*=0.002, *d*=1.01) and low activity (*P*=0.010, *d*=0.68) groups. In the male low-activity group, Arabian jasmine also had an effect (*P<*0.001, *d*=0.50), causing a shift to higher distance from the bottom on the second day. The vehicle treated high-activity group had a trend towards changing to a higher distance from the bottom (*P*=0.070).

For the center distance of female AB zebrafish, we found no effect of Arabian jasmine (*P*=0.35), activity (*P*=0.21), and no interaction (*P*=0.21). For male AB zebrafish, we found a medium-sized effect of Arabian jasmine (*P*=0.016, η^2^=0.08), but no effect of activity (*P*=0.34), or an interaction (*P*=0.61). Post-hoc tests found that Arabian jasmine treated low-activity males swam closer to the center of the tank compared to the vehicle group (*P*=0.046, *d*=0.69). For the change in center distance over time, there was an effect in Arabian jasmine treated high-activity females (*P*=0.02, *d*=0.46) and vehicle treated low-activity males (*P*=0.02, *d*=0.86) where they were both closer to the border of the tank. There was also a trend towards an effect in Arabian jasmine treated high-activity males (*P*=0.06, *d*=0.53).

For the percentage of exploration in female AB zebrafish, we found a medium-sized effect of Arabian jasmine (*P*=0.040, η^2^=0.06), but no effect of activity (*P*=0.83), or an interaction (*P*=0.61). Post-hoc tests found that the Arabian jasmine treated low-activity females had a trend toward a medium-sized effect of exploring the tank more than the vehicle group (*P*=0.091, *d*=0.57). For the percentage of exploration of male AB zebrafish, we found no effect of Arabian jasmine (*P*=0.40), activity (*P*=0.12), and no interaction (*P*=0.86). For the change in percentage of exploration over time, we found no change in exploration behavior in either vehicle or Arabian jasmine groups.

For the distance travelled of female AB zebrafish, we found a medium-sized effect of activity (*P*=0.004, η^2^=0.12), but no effect of Arabian jasmine (*P*=0.64), or interaction (*P*=0.91). For the distance travelled of male AB zebrafish, we found a medium-sized effect of activity (*P*=0.011, η^2^=0.09), but no effect of Arabian jasmine (*P*=0.29), or an interaction (*P*=0.37). However, post-hoc tests found no differences for either females or males. For the change in distance travelled over time, we found an effect in vehicle (*P*=0.044, *d*=0.72) and Arabian jasmine (*P*=0.003, *d*=0.89) treated low-activity females, both of which had more activity than on day 1. The vehicle treated high-activity females (*P*=0.090, *d*=0.56) and Arabian jasmine treated low-activity males (*P*=0.080, *d*=0.51) had trends toward less activity than day 1. Arabian jasmine treated high-activity males also had a trend toward less activity (*P*=0.090, *d*=0.51).

## Discussion

We find that sex, genetic background, and personality all contribute to the anxiety-related behavioral effects of Arabian jasmine in adult zebrafish. This conclusion is based on the following findings: (1) Arabian jasmine caused a decrease in bottom distance, and to a lesser extent percent explored, (i.e., was anxiogenic) in female AB fish (Fig. 2A); there was no effect in AB males, and no effect in WIK or TL fish of either sex. (2) Arabian jasmine increased percent explored, and to a lesser extent bottom distance (i.e., was anxiolytic) in shy, but not bold, female AB fish (Fig. 3D), and decreased center distance in male AB fish with low activity (Fig. 3F). Taken together, these data suggest that strain, sex, and personality interact to modulate the effects of Arabian jasmine on behavior.

Our paper is the first to find that Arabian jasmine affects behavior in zebrafish. Given that Arabian jasmine is a complex mixture of many phytochemicals, it is not clear what constituents of Arabian jasmine may be driving these behavioral effects. Chemical analysis of our Arabian jasmine preparation may provide some clues. Our preparation had particularly high concentrations of linalool (28.4%) and dimethyl sulfide (30.3 %). We also isolated lesser amounts of other potentially psychoactive compounds such as 3-hexen-1-ol (0.5%), linalool oxide (0.09%), phenylethyl alcohol (1.46%) and benzaldehyde (0.68%). Most of these compounds have been described as anxiolytic [3,9,10,29–34] except for benzaldehyde [11]. However, there are no reports on the behavioral effects of dimethyl sulfide, which is known for having an unpleasant odor [35].

The high concentration of linalool in our extract is notable because it has been consistently found to be anxiolytic [10,30–32,36]. This may be via the many effects linalool has been found to have on neurotransmission. For example, linalool enhances GABAergic currents in mice [36] and directly inhibits glutamatergic neurons [37,38]. Linalool has also been found to modulate serotonergic functioning, where Jarvis et al. [31] found that linalool acts as a 5-HT_3_ antagonist. 5-HT₃ receptors are ionotropic receptors that mediate excitatory neurotransmission; their activity has been linked to anxiety [39,40]. Linalool affects other neurotransmitter systems as well, like dopamine and norepinephrine [30]. At the physiological level, Höferl et al. [32] found linalool to promote relaxation in humans alongside a reduction in cortisol and heart rate, and Yamamoto et al. [10] found that inhaled linalool in restrained mice had reduced hypothalamic-pituitary-adrenal (HPA) activity. Behaviorally, linalool was found to be anxiolytic in mice [30,41] and to cause a reduction in aggression in male mice [9]. Taken together with the present study, these findings suggest that the anxiolytic effects of Arabian jasmine may be mediated by linalool’s effects on neurotransmitters and stress hormone activity in zebrafish.

In contrast to linalool, benzaldehyde is one of the few compounds in Arabian jasmine that has been described as anxiogenic, although the evidence here is weaker as there is only one study describing its effects. Laham et al. [11] found that inhalation of benzaldehyde by Sprague-Dawley rats resulted in increased aggressive behavior, noise sensitivity, and reduced motor activity. Interestingly, Tabatabaie and Floyd [42] found that benzaldehyde inactivates the antioxidant enzyme glutathione peroxidase, which can disrupt the redox balance; this may lead to elevated anxiety via an increase in oxidative stress [43]. Thus, it may be the case that the anxiogenic effects we observed in female AB zebrafish are due to higher sensitivity of these fish to benzaldehyde. However, given the complex mixture of chemicals in Arabian jasmine, more research would be needed to more strongly draw this conclusion.

There are some notable differences in the chemicals we identified in Arabian jasmine compared to previous reports. For example, we find higher linalool (28.4%) than prior work (2-10%) and dimethyl sulfide has not previously been reported to be a constituent of *J. sambac* [7,8,44]. These differences may be because we used ultra-sonic assisted extraction where microbubbles break down plant walls [45]. This contrasts with traditional methods that use warm water [1] that can degrade volatile compounds [46]. Additionally, our air pressure-assisted extraction may enhance this process by improving mass transfer efficiency [47]. Thus, compared with traditional methods, our approach minimizes the loss of volatile compounds. Different preparation methods are a challenge for the field of traditional medicine as it has been found that the intensity, solvent choice, and temperature used in ultrasonic-assisted extraction can cause the breakdown of active compounds [45]. For example, the high concentration of dimethyl sulfide we observed may have been produced through a Maillard reaction between glucose and methionine in the plant, with ultrasonic treatment accelerating the degradation of methionine and promoting the formation of this volatile sulfur compound [48,49]. More work is clearly needed to better understand the impact of extraction techniques on the phytochemical profile and subsequent behavioral effects of Arabian jasmine.

We found several instances in which genetic background and sex influence the effects of Arabian jasmine on behavior. Arabian jasmine caused female AB zebrafish to decrease their bottom distance and exploratory behavior, while there was no effect in males, and no effect in WIK or TL fish of either sex. Genetic variation is well known to influence a wide variety of behavioral responses [26,27,50,51]. This is likely due, in part, to differences in neurochemical profiles [52–54] that may affect drug responses. For example, strain specific effects of anxiolytics, like diazepam and losartan, have been observed in mice where diazepam induced anxiolytic effects in C57BL/6, DBA/2, and BKW mice whereas losartan produced effects only in BKW mice [55]. Similarly, Dlugos and Robin [56] found that strain affected behavioral responses to ethanol in in WT, LFS and BLF zebrafish where only they observed anxiolytic effects in the WT and LFS strains. Beyond strain differences, sex also plays a significant role in drug responses. Male and female zebrafish differ in their responses to ethanol, where females are more sensitive than males [12,57]. Johnson et al. [58] also investigated the effect of chondroitin sulfate on swimming velocity in zebrafish, finding increased velocity only in males. There are also several baseline differences in behavior between the sexes. For example, Fontana et al. [59] identified sex differences in anxiety-related behaviors in male and female zebrafish. And Rajput et al. [27] used a three-dimensional novel tank test to observe sex-based differences in anxiety-related behaviors, including distance from the center, distance traveled, and percentage of exploration. Sex differences are also seen at the neurochemical level in zebrafish where Beigloo et al. [15] found that males have higher dopamine and serotonin than females. Thus, the differential responses to Arabian jasmine observed in this study is likely driven by an interaction of genetics and sex, possibly due to variations in in neurotransmitter systems.

We found that different behavioral types (i.e., personality) in zebrafish altered the anxiety-related response of Arabian jasmine. For example, we found Arabian jasmine increased exploratory behaviors in shy females and weakly elevated bottom distance in low-activity males. Variations in drug responses based on personality types have been observed in mice and zebrafish in a handful of cases. For instance, Beigloo et al. [15] found differences in the responses of bold and shy WIK zebrafish following escitalopram administration. In mice, Mehrhoff et al. [16] found that diazepam had an anxiolytic effect only low-activity mice. One possible explanation for these effects is differences in gene expression that may underlie behavioral types. For example, Norton et al. [60] identified *fgfr1a* as a key gene influencing boldness behavior in zebrafish. Similarly, Thomas et al. [61] studied genetic variation in inbred mice with high and low open-field activity and identified *Ctsc* and *Vmn1r1* as genes linked to differing activity levels. Finally, Booher et al. [62] showed that *Ndufa13* gene encoding the supernumerary subunit A13 of nicotinamide adenine dinucleotide dehydrogenase complex I, is a key candidate gene influencing high or low activity in mice. Thus, the varied responses to Arabian jasmine based on personality-type in zebrafish we observed may be driven by associated interindividual differences in gene expression.

In this study, we observed conflicting results: the 10 mg kg⁻¹ Arabian jasmine treatment group in female AB zebrafish exhibited more anxiety-like behavior, while Arabian jasmine had an anxiolytic effect in shy female AB zebrafish. This phenomenon, known as a paradoxical drug effect, has been reported with anxiolytic drugs such as benzodiazepines [63] and fluoxetine [64]. For example, in patients who have genetic links to psychological disorders, anxiogenic effects of benzodiazepines were more likely [63]. In addition, the age of mice can influence responses to fluoxetine, where adults show anxiolytic effect and, juveniles an anxiogenic effect [64]. One possible explanation for this paradoxical effect could be differences in HPA axis function. Lindholm et al. [65] found that genetic variation can influence the functioning of the HPA axis and associated neurochemicals, potentially contributing to paradoxical drug responses. Beigloo et al. [15] also found the differences in serotonergic system can affect the boldness of female zebrafish, which could help explain these unexpected drug responses. In mice, Krakenberg et al. [66] showed that variations in the 5-HT genotype (e.g., serotonin transporter (5-HTT) gene) were associated with different levels of anxiety-related behavior in mice. The level of GABA may also play an important role in paradoxical drug effects. Majewska [67] found that variation in neuro-steroid levels can influence GABA_A_ receptor activity. These findings suggest that differing neurochemical profiles in fish of different background strains or different personality types may contribute to the paradoxical response to Arabian jasmine in zebrafish.

This study has limitations and should be mentioned. For example, we administered Arabian jasmine acutely, with a single dose, whereas humans typically consume it chronically [1]. Additionally, humans often use herbal preparations containing a combination of multiple plants. This study selected a dose of Arabian jasmine based on its use in such preparations; however, for simplicity of interpretation, the effects of interactions with other plants were not examined. We also did not examine biochemical correlates of anxiety, like HPA axis function. Future work should explore cortisol level to identify potential differences in HPA axis functioning across strain, sex, and personality that could contribute to the variable effects of Arabian jasmine [68]. Another factor that we did not examine here that could influence the effects of Arabian jasmine are differences in pharmacokinetic profiles across strains and sexes. Finally, it is difficult to extrapolate the anxiety-like behavior observed here to the phenomenological experience of humans taking Arabian jasmine. Nonetheless, we believe our work lays the foundation for using zebrafish to unravel the biological basis for the complex behavioral effects of Arabian jasmine and other natural compounds used in traditional medicine.

## Conclusion

In summary, our work reveals that the effects of Arabian jasmine on anxiety-related behavior is affected by background genetics, sex and personality. Further observations are needed to explore the mechanism of action based on the molecular and neural basis for these different responses. However, our results lead to the conclusion that the paradoxical responses of Arabian jasmine that have previously been reported may be due to differences in the characteristics of subjects.

## Methods

### Plant materials

*Jasminum sambac* (L.) Aiton (Fig. 1A) was collected from Arabian jasmine cultivation in Khumphaeng Phet, Thailand (16°27’45.0"N 99°40’11.3"E) on the morning (8-10AM) of 25^th^ April 2023. Voucher specimens (herbarium number: PBM006430) were made using standard herbarium specimen preparation procedures [69] and deposited in the herbarium at Sireeruckhachati Nature Learning Park, Mahidol University, Thailand. The Flora of China [70] and the Flora of Thailand [71], as well as a recent document of the genus *Jasminum* in Thailand [72], were used for identification.

### Plant extraction and chemical identification

To replicate the preparation used in traditional medicine, water was chosen as the solvent. This process was followed by ultrasonic- and pressure-assisted extraction methods (Fig. 1B). The ultrasonic-assisted extraction method was adapted from Vinatoru et al. [73], while the pressure-assisted extraction was modified from Ong et al. [74]. Arabian jasmine extract was prepared by sonicating Arabian jasmine flowers in water at a ratio of 5 parts Arabian jasmine to 3 parts water by weight for 15 minutes. The resulting aqueous solution was then subjected to continuous extraction under reduced air pressure (400 mmHg) using a rotary evaporator for 30 minutes.

After filtration, the extract was stored in an airtight, light-protected container at -4 °C. To create a concentrated solution, 100 mL of the Arabian jasmine extract was evaporated using a rotary evaporator to yield a concentrate of 25 mg mL⁻¹.

Following the extraction, a gas chromatograph equipped with a headspace extractor (7697A Static Headspace Sampler and Agilent 7890B System, Agilent Technologies, Santa Clara, CA, USA) was used for chemical analysis. The system included an HP-INNOWax capillary column (30 m length × 250 μm diameter × 0.25 μm thickness, Agilent Technologies) and a mass spectrometer with an ion trap detector (Agilent Technologies). The analysis was performed in the range of 33 to 400 m/z under ionization energy of 70 eV. n-Alkane compounds were used as external reference standards to ensure accuracy.

The chromatographic analysis of the Arabian jasmine extract was conducted by comparing retention times, peak areas, peak heights, and mass spectral patterns with those in the NIST library [75] of authentic compounds.

### Gelatin preparation and gelatin-feed habituation

For gelatin feed preparation, we followed the method from Ochocki and Kenney [76]. Arabian jasmine water extract was administrated using a gelatin-based feed at dose of 5, 10, or 20 mg kg^-1^ body mass. This was determined to approximately match the suggested doses in traditional preparations given to people [1]. The gelatin feed consisted of 12% w/v gelatin (Sigma-Aldrich), 4% w/v spirulina (Argent Aquaculture) and brine shrimp extract. The extract was made by suspending 250 mg ml^−1^ of micro fine brine shrimp (Brine Shrimp Direct) in water followed by 1 h of stirring. The suspension was then centrifuged twice at 12,500 *g*, keeping the supernatant each time, and then diluted in two volumes of water before addition to the gelatin feed mixture. Arabian jasmine and water were added before warming the solution to 45°C. After warming, drug- or vehicle-containing solution was pipetted into individually sized morsels for feeding at 1% body mass. Gelatin was allowed to set on ice for at least 20 min before feeding.

Fish were individually habituated to the gelatin-feed for three days before behavior tests. On each day of the experiment, individuals were removed from the housing racks to the behavioral room for 1 h before being given a non-dosed gelatin feed instead of their morning feed. Individuals were isolated by the placement of transparent barriers in the tanks for 1 h. Once feed was given, we recorded whether individuals ate the feed within 5 min.

### Subjects

Subjects were female and male zebrafish of the AB, TL and WIK strains. Fish were 4–6 months of age and raised at Wayne State University. Housing was in high-density racks under standard conditions (water temperature: 27.5±0.5°C, salinity: 500±10 µS and pH 7.4±0.2) with a 14 h:10 h light: dark cycle (lights on at 08:00 h). Fish were fed twice daily, in the morning with a dry feed (Gemma 300, Skretting) and in the afternoon with brine shrimp (*Artemia salina*, Brine Shrimp Direct). The sex of fish was determined using three secondary sex characteristics: shape, color and presence of pectoral fin tubercles [77]. Sex was confirmed following euthanasia by the presence or absence of eggs. All procedures were approved by the Wayne State University Institutional Animal Care and Use Committee (Protocol ID: 21-02-3238).

### Novel tank and mirror biting tests

We used the novel tank test (NTT) to examine anxiety-related behavior, following the method described by Rajput et al. [27]. Additionally, the mirror biting test (MBT) was employed to assess aggressive behavior.

One week before the experiment, fish were housed in 2 L tanks, with two female/male pairs per tank. Each tank was divided in half using a transparent divider, allowing one pair per side. To acclimate to the behavioral testing environment, tanks were removed from their housing racks and transferred to the behavioral room. Three days before Arabian jasmine administration, fish were acclimated to the gelatin feed to ensure familiarity with the feeding method. Behavioral testing was conducted between 09:00 h and 14:00 h, with fish placement counterbalanced across sex and drug treatment groups. To avoid the buildup of chemical cues released by the fish during testing, NTT water was replaced between animals.

For the NTT, experimental tanks were five-sided (15×15×15 cm) and made from frosted acrylic (TAP Plastics). Each tank was filled to a height of 12 cm with 2.5 L of fish facility water and placed in a white Plasticore enclosure to diffuse light and minimize external disturbances. Fish were individually placed into the tanks for 6 minutes while video recordings were made. Fish were tracked in the videos using DeepLabCut [78].

Following the NTT, fish were transferred to MBT tanks for another 6-minute session, during which video recordings were also made. The MBT tanks were identical to the NTT tanks, except for the inclusion of a mirror on the right-side wall. Like the NTT tanks, they were filled with 2.5 L of fish facility water and housed in a white Plasticore enclosure. D435 Intel RealSense™ depth-sensing cameras (Intel) were mounted 20 cm above the tanks to record the sessions. These cameras were connected to Linux workstations via high-speed USB cables (NTC Distributing). The MBT videos were first tracked using DeepLabCut [78] and then SimBA [79] was used to quantify social and aggressive behavior. We followed the behavioral terms of Blaser and Gerlai [80] (attack time as aggressive behavior) and Porfiri et al. [81] (parallel swimming time as social behavior). To measure attack time, we calculated the total time a fish spent attacking within 3 cm of the mirror (discrimination threshold: 0.35; minimum bout length: 20 ms). Similarly, to measure parallel swimming, we calculated the total time a fish spent swimming parallel to its reflection within 3 cm of the mirror (discrimination threshold: 0.4; minimum bout length: 50 ms).

### Dose response and genetic variation

After habituation to the gelatin-feed, individuals were given either a vehicle or Arabian jasmine containing feed 30 min before being placed in the experimental tank. Only fish that ate the feed within 5 min were included in the analysis (3 female and 7 male fish given vehicle were excluded). We centered our dose response around the 10 mg kg^-1^ dose based on the traditional preparation used for people [1]. Thirty minutes was chosen because several studies in zebrafish have found that oral drug administration results in peak serum concentrations within 40 min [82,83], and previous studies found behavioral response of drug administration using this method at this time point [76].

### Identifying personality type

For the effect of Arabian jasmine on the personality of zebrafish experiment, we followed the method of Beigloo et al. [15]. The behavior tests occurred over 2 days. On day 1, all individuals were given non-dosed feed 30 min before being placed in the novel tank. The boldness (z-scores of bottom distance and the percentage of exploration added together) and activity (z-score of distance travelled) categories were calculated from day 1. On day 2 of the anxiety-related tests, individuals were given either vehicle or Arabian jasmine containing feed 30 min before being placed in the tanks. The AB zebrafish and dose of 10 mg kg^-1^ Arabian jasmine were selected based on the results of the dose-response experiment.

### Data analysis and visualization

Data analysis was performed using R version 4.4.0 [84]. Graphs were made using ggplot2 [85]. Statistical analysis was done using a 4 × 2 (treatment × sex) ANOVA, a 2 × 2 (treatment × boldness/activity) ANOVA, or a one-sample *t*-test (the change of behavior over time). All tests were two-tailed. Omnibus tests were followed up with Dunnett’s multiple comparison tests to examine the effects of Arabian jasmine dosage [86] and pairwise *t*-test with False Discovery Rate (FDR) correction to compare the effects Arabian jasmine within sex [87]. For effect sizes, ANOVAs are reported as η^2^ and t-tests as Cohen’s d. The interpretation of effect sizes as small (0.01<η^2^<0.06; 0.2<d<0.5), medium (0.06≤η^2^<0.14; 0.5≤d<0.8) or large (η^2^≥0.14; d≥0.8) [88].

## Supporting information

Figure S1 Influence of Arabian jasmine on aggressive behaviors based on personality.

Table S1 GC-MS profiling of Arabian jasmine water extract.

Table S2 novel tank test dose response data.

Table S3 mirror biting test dose response data.

Table S4 personality data.

Table S5 novel tank test personality data.

Table S6 mirror biting test personality data.

## Data availability

Relevant data can be found within the article and its supplementary information. Data and supplementary information are available from Figshare: 10.6084/m9.figshare.28295993.

## Acknowledgments and Funding

Funding was provided by NIGMS (R35GM142566 to JWK) and the Science Achievement Scholarship of Thailand (SAST to TA and NP). The funders had no role in study design, data collection and analysis, decision to publish, or preparation of the manuscript. We thank Dinh Luong for the excellent maintenance and husbandry of our zebrafish colony.

## Notes

### Competing Interest Statement

The authors have declared no competing interest.

## References

1. Ministry of Public Health. National Thai Traditional Medicine Formulary 2021 Edition. Bangkok, Thailand: Samchareon phanich; 2021.

2. Kafaei M, Burry J, Latifi M, Ciorciari J, Aminitabar A. Investigating the effect of different smells in the indoor built environment (office) on human emotions using a mixed-method approach. Archit Sci Rev. 2024;67: 518–528. doi:10.1080/00038628.2024.2370435

3. Kuroda K, Inoue N, Ito Y, Kubota K, Sugimoto A, Kakuda T, et al. Sedative effects of the jasmine tea odor and (R)-(−)-linalool, one of its major odor components, on autonomic nerve activity and mood states. Eur J Appl Physiol. 2005;95: 107–114. doi:10.1007/s00421-005-1402-8

4. Yadegari M, Mahmoodi-Shan GR, Kamkar MZ, Vakili M. Effects of inhaling jasmine essential oil on anxiety and blood cortisol levels in candidates for laparotomy: A randomized clinical trial. J Nurs Midwifery Sci. 2021;8: 128–133. doi:10.4103/jnms.jnms_125_20

5. Hongratanaworakit T. Stimulating effect of aromatherapy massage with jasmine oil. Nat Prod Commun. 2010;5: 157–162.

6. Sayowan W, Siripornpanich V, Hongratanaworakit T, Kotchabhakdi N, Ruangrungsi N. The Effects of Jasmine Oil Inhalation on Brain Wave Activies and Emotions. J Health Res. 2013;27: 73–77.

7. Khidzir KM, Cheng SF, Chuah CH. Interspecies variation of chemical constituents and antioxidant capacity of extracts from Jasminum sambac and Jasminum multiflorum grown in Malaysia. Ind Crops Prod. 2015;74: 635–641. doi:10.1016/J.INDCROP.2015.05.053

8. Younis A, Mehdi A, Riaz A. Supercritical carbon dioxide extraction and gas chromatography analysis of Jasminum sambac essential oil. Pak J Bot. 2011;43: 163–168.

9. Linck VM, da Silva AL, Figueiró M, Caramão EB, Moreno PRH, Elisabetsky E. Effects of inhaled Linalool in anxiety, social interaction and aggressive behavior in mice. Phytomedicine. 2010;17: 679–683. doi:10.1016/j.phymed.2009.10.002

10. Yamamoto N, Fujiwara S, Saito-iizumi K, Kamei A, Shinozaki F, WATANABE Y, et al. Effects of Inhaled (S)-Linalool on Hypothalamic Gene Expression in Rats under Restraint Stress. Biosci Biotechnol Biochem. 2013;77: 2413–2418. doi:10.1271/bbb.130524

11. Laham S, Broxup B, Robinet M, Potvin M, Schrader K. Subacute inhalation toxicity of benzaldehyde in the Sprague-Dawley rat. Am Ind Hyg Assoc J. 1991;52: 503–510. doi:10.1080/15298669191365126

12. Dlugos CA, Brown SJ, Rabin RA. Gender differences in ethanol-induced behavioral sensitivity in zebrafish. Alcohol. 2011;45: 11–18. doi:10.1016/j.alcohol.2010.08.018

13. Hilderbrand ER, Lasek AW. Studying Sex Differences in Animal Models of Addiction: An Emphasis on Alcohol-Related Behaviors. ACS Chem Neurosci. 2018;9: 1907–1916. doi:10.1021/acschemneuro.7b00449

14. Pirmohamed M. Pharmacogenomics: current status and future perspectives. Nat Rev Genet. 2023;24: 350–362. doi:10.1038/s41576-022-00572-8

15. Beigloo F, Davidson CJ, Gjonaj J, Perrine SA, Kenney JW. Individual differences in the boldness of female zebrafish are associated with alterations in serotonin function. J Exp Biol. 2024;227: jeb247483. doi:10.1242/jeb.247483

16. Mehrhoff EA, Booher WC, Hutchinson J, Schumacher G, Borski C, Lowry CA, et al. Diazepam effects on anxiety-related defensive behavior of male and female high and low open-field activity inbred mouse strains. Physiol Behav. 2023;271: 114343. doi:10.1016/j.physbeh.2023.114343

17. Parichy DM. Advancing biology through a deeper understanding of zebrafish ecology and evolution. eLife. 2015;4. doi:10.7554/eLife.05635

18. Burgess HA, Burton EA. A Critical Review of Zebrafish Neurological Disease Models−1. The Premise: Neuroanatomical, Cellular and Genetic Homology and Experimental Tractability. Oxf Open Neurosci. 2023;2: kvac018. doi:10.1093/oons/kvac018

19. Gerlai R. Zebrafish (Danio rerio): A newcomer with great promise in behavioral neuroscience. Neurosci Biobehav Rev. 2023;144: 104978. doi:10.1016/j.neubiorev.2022.104978

20. Kenney JW. Associative and nonassociative learning in adult zebrafish. Behavioral and Neural Genetics of Zebrafish. Elsevier; 2020. pp. 187–204. doi:10.1016/b978-0-12-817528-6.00012-7

21. Howe K, Clark MD, Torroja CF, Torrance J, Berthelot C, Muffato M, et al. The zebrafish reference genome sequence and its relationship to the human genome. Nature. 2013;496: 498–503. doi:10.1038/nature12111

22. Panula P, Sallinen V, Sundvik M, Kolehmainen J, Torkko V, Tiittula A, et al. Modulatory neurotransmitter systems and behavior: towards zebrafish models of neurodegenerative diseases. Zebrafish. 2006;3: 235–247. doi:10.1089/zeb.2006.3.235

23. Panula P, Chen YC, Priyadarshini M, Kudo H, Semenova S, Sundvik M, et al. The comparative neuroanatomy and neurochemistry of zebrafish CNS systems of relevance to human neuropsychiatric diseases. Neurobiol Dis. 2010;40: 46–57. doi:10.1016/j.nbd.2010.05.010

24. Alsop D, Vijayan MM. Development of the corticosteroid stress axis and receptor expression in zebrafish. Am J Physiol Regul Integr Comp Physiol. 2008;294. doi:10.1152/ajpregu.00671.2007

25. Kalueff AV, Echevarria DJ, Stewart AM. Gaining translational momentum: more zebrafish models for neuroscience research. Prog Neuropsychopharmacol Biol Psychiatry. 2014;55: 1–6. doi:10.1016/j.pnpbp.2014.01.022

26. Audira G, Siregar P, Strungaru S-A, Huang J-C, Hsiao C-D. Which Zebrafish Strains Are More Suitable to Perform Behavioral Studies? A Comprehensive Comparison by Phenomic Approach. Biology. 2020;9: 200. doi:10.3390/biology9080200

27. Rajput N, Parikh K, Kenney JW. Beyond bold versus shy: Zebrafish exploratory behavior falls into several behavioral clusters and is influenced by strain and sex. Biol Open. 2022;11: bio059443. doi:10.1242/bio.059443

28. Tran S, Gerlai R. Individual differences in activity levels in zebrafish (*Danio rerio*). Behav Brain Res. 2013;257: 224–229. doi:10.1016/j.bbr.2013.09.040

29. Tokumo K, Tamura N, Hirai T, Nishio H. Effects of (*Z*)-3-hexenol, a major component of green odor, on anxiety-related behavior of the mouse in an elevated plus-maze test and biogenic amines and their metabolites in the brain. Behav Brain Res. 2006;166: 247–252. doi:10.1016/j.bbr.2005.08.008

30. Cheng B-H, Sheen L-Y, Chang S-T. Evaluation of anxiolytic potency of essential oil and *S*-(+)-linalool from *Cinnamomum osmophloeum* ct. linalool leaves in mice. J Tradit Complement Med. 2015;5: 27–34. doi:10.1016/j.jtcme.2014.10.007

31. Jarvis GE, Barbosa R, Thompson AJ. Noncompetitive Inhibition of 5-HT3 Receptors by Citral, Linalool, and Eucalyptol Revealed by Nonlinear Mixed-Effects Modeling. J Pharmacol Exp Ther. 2016;356: 549–562. doi:10.1124/jpet.115.230011

32. Höferl M, Krist S, Buchbauer G. Chirality Influences the Effects of Linalool on Physiological Parameters of Stress. Planta Med. 2006;72: 1188–1192. doi:10.1055/s-2006-947202

33. Souto-Maior FN, Carvalho FL de, Morais LCSL de, Netto SM, de Sousa DP, Almeida RN de. Anxiolytic-like effects of inhaled linalool oxide in experimental mouse anxiety models. Pharmacol Biochem Behav. 2011;100: 259–263. doi:10.1016/j.pbb.2011.08.029

34. Ramadan B, Cabeza L, Cramoisy S, Houdayer C, Andrieu P, Millot J-L, et al. Beneficial effects of prolonged 2-phenylethyl alcohol inhalation on chronic distress-induced anxio-depressive-like phenotype in female mice. Biomed Pharmacother. 2022;151: 113100. doi:10.1016/j.biopha.2022.113100

35. Tonzetich J. Production and Origin of Oral Malodor: A Review of Mechanisms and Methods of Analysis. J Periodontol. 1977;48: 13–20. doi:10.1902/jop.1977.48.1.13

36. Milanos S, Elsharif SA, Janzen D, Buettner A, Villmann C. Metabolic Products of Linalool and Modulation of GABAA Receptors. Front Chem. 2017;5. doi:10.3389/fchem.2017.00046

37. Batista PA, Werner MF d. P, Oliveira EC, Burgos L, Pereira P, Brum LF d. S, et al. Evidence for the involvement of ionotropic glutamatergic receptors on the antinociceptive effect of (-)-linalool in mice. Neurosci Lett. 2008;440: 299–303. doi:10.1016/j.neulet.2008.05.092

38. Silva Brum LF, Emanuelli T, Souza DO, Elisabetsky E. Effects of Linalool on Glutamate Release and Uptake in Mouse Cortical Synaptosomes. Neurochem Res. 2001;26: 191–194. doi:10.1023/A:1010904214482

39. Costall B, Naylor RJ, Tyers MB. The psychopharmacology of 5-HT3 receptors. Pharmacol Ther. 1990;47: 181–202. doi:10.1016/0163-7258(90)90086-H

40. Du Y, Li Z, Zhao Y, Han J, Hu W, Liu Z. Role of 5-hydroxytryptamine type 3 receptors in the regulation of anxiety reactions. J Zhejiang Univ-Sci B. 2024;25: 23–37. doi:10.1631/jzus.B2200642

41. Harada H, Kashiwadani H, Kanmura Y, Kuwaki T. Linalool Odor-Induced Anxiolytic Effects in Mice. Front Behav Neurosci. 2018;12. doi:10.3389/fnbeh.2018.00241

42. Tabatabaie T, Floyd RA. Inactivation of Glutathione Peroxidase by Benzaldehyde. Toxicol Appl Pharmacol. 1996;141: 389–393. doi:10.1006/taap.1996.0304

43. Schriever SC, Zimprich A, Pfuhlmann K, Baumann P, Giesert F, Klaus V, et al. Alterations in neuronal control of body weight and anxiety behavior by glutathione peroxidase 4 deficiency. Neuroscience. 2017;357: 241–254. doi:10.1016/j.neuroscience.2017.05.050

44. Zhang J, Li J, Wang J, Sun B, Liu Y, Huang M. Characterization of aroma-active compounds in Jasminum sambac concrete by aroma extract dilution analysis and odour activity value. Flavour Fragr J. 2021;36: 197–206. doi:10.1002/FFJ.3631

45. Kumar K, Srivastav S, Sharanagat VS. Ultrasound assisted extraction (UAE) of bioactive compounds from fruit and vegetable processing by-products: A review. Ultrason Sonochem. 2020;70: 105325. doi:10.1016/j.ultsonch.2020.105325

46. Santos MB, Sillero L, Gatto DA, Labidi J. Bioactive molecules in wood extractives: Methods of extraction and separation, a review. Ind Crops Prod. 2022;186: 115231. doi:10.1016/j.indcrop.2022.115231

47. Huang H-W, Cheng M-C, Chen B-Y, Wang C-Y. Effects of high pressure extraction on the extraction yield, phenolic compounds, antioxidant and anti-tyrosinase activity of Djulis hull. J Food Sci Technol. 2019;56: 4016–4024. doi:10.1007/s13197-019-03870-y

48. Yu H, Keh MZM, Seow Y-X, Ong PKC, Zhou W. Kinetic Study of High-Intensity Ultrasound-Assisted Maillard Reaction in a Model System of D-Glucose and L-Methionine. Food Bioprocess Technol. 2017;10: 1984–1996. doi:10.1007/s11947-017-1971-7

49. Yu T-H, Ho C-T. Volatile Compounds Generated from Thermal Reaction of Methionine and Methionine Sulfoxide with or without Glucose. J Agric Food Chem. 1995;43: 1641–1646. doi:10.1021/jf00054a043

50. Burne T, Scott E, van Swinderen B, Hilliard M, Reinhard J, Claudianos C, et al. Big ideas for small brains: what can psychiatry learn from worms, flies, bees and fish? Mol Psychiatry. 2011;16: 7–16. doi:10.1038/mp.2010.35

51. Crawley JN, Belknap JK, Collins A, Crabbe JC, Frankel W, Henderson N, et al. Behavioral phenotypes of inbred mouse strains: implications and recommendations for molecular studies. Psychopharmacol Berl. 1997;132: 107–24.

52. Pan Y, Chatterjee D, Gerlai R. Strain dependent gene expression and neurochemical levels in the brain of zebrafish: Focus on a few alcohol related targets. Physiol Behav. 2012;107: 773–780. doi:10.1016/j.physbeh.2012.01.017

53. Waller SB, Ingram DK, Reynolds MA, London ED. Age and Strain Comparisons of Neurotransmitter Synthetic Enzyme Activities in the Mouse. J Neurochem. 1983;41: 1421–1428. doi:10.1111/j.1471-4159.1983.tb00841.x

54. Tagawa N, Sugimoto Y, Yamada J, Kobayashi Y. Strain differences of neurosteroid levels in mouse brain. Steroids. 2006;71: 776–784. doi:10.1016/j.steroids.2006.05.008

55. Gard PR, Haigh SJ, Cambursano PT, Warrington CA. Strain differences in the anxiolytic effects of losartan in the mouse. Pharmacol Biochem Behav. 2001;69: 35–40. doi:10.1016/S0091-3057(01)00491-9

56. Dlugos CA, Rabin RA. Ethanol effects on three strains of zebrafish: model system for genetic investigations. Pharmacol Biochem Behav. 2003;74: 471–480. doi:10.1016/S0091-3057(02)01026-2

57. Souza TP, Franscescon F, Stefanello FV, Müller TE, Santos LW, Rosemberg DB. Acute effects of ethanol on behavioral responses of male and female zebrafish in the open field test with the influence of a non-familiar object. Behav Processes. 2021;191: 104474. doi:10.1016/j.beproc.2021.104474

58. Johnson AL, Hurd PL, Hamilton TJ. Sex, drugs, and zebrafish: Acute exposure to anxiety-modulating compounds in a modified novel tank dive test. Pharmacol Biochem Behav. 2024;243: 173841. doi:10.1016/j.pbb.2024.173841

59. Fontana BD, Cleal M, Parker MO. Female adult zebrafish (Danio rerio) show higher levels of anxiety-like behavior than males, but do not differ in learning and memory capacity. Eur J Neurosci. 2020;52: 2604–2613. doi:10.1111/ejn.14588

60. Norton WHJ, Stumpenhorst K, Faus-Kessler T, Folchert A, Rohner N, Harris MP, et al. Modulation of Fgfr1a Signaling in Zebrafish Reveals a Genetic Basis for the Aggression–Boldness Syndrome. J Neurosci. 2011;31: 13796–13807. doi:10.1523/JNEUROSCI.2892-11.2011

61. Thomas AL, Evans LM, Nelsen MD, Chesler EJ, Powers MS, Booher WC, et al. Whole-Genome Sequencing of Inbred Mouse Strains Selected for High and Low Open-Field Activity. Behav Genet. 2021;51: 68–81. doi:10.1007/s10519-020-10014-y

62. Booher WC, Vanderlinden LA, Hall LA, Thomas AL, Evans LM, Saba LM, et al. Hippocampal RNA sequencing in mice selectively bred for high and low activity. Genes Brain Behav. 2023;22: e12832. doi:10.1111/gbb.12832

63. Gutierrez MA, Roper JM, Hahn P. Paradoxical Reactions to Benzodiazepines: When to expect the unexpected. AJN Am J Nurs. 2001;101: 34.

64. Oh J, Zupan B, Gross S, Toth M. Paradoxical Anxiogenic Response of Juvenile Mice to Fluoxetine. Neuropsychopharmacology. 2009;34: 2197–2207. doi:10.1038/npp.2009.47

65. Lindholm H, Morrison I, Krettek A, Malm D, Novembre G, Handlin L. Genetic risk-factors for anxiety in healthy individuals: polymorphisms in genes important for the HPA axis. BMC Med Genet. 2020;21: 184. doi:10.1186/s12881-020-01123-w

66. Krakenberg V, von Kortzfleisch VT, Kaiser S, Sachser N, Richter SH. Differential Effects of Serotonin Transporter Genotype on Anxiety-Like Behavior and Cognitive Judgment Bias in Mice. Front Behav Neurosci. 2019;13. doi:10.3389/fnbeh.2019.00263

67. Majewska MD. Neurosteroids: Endogenous bimodal modulators of the GABAA receptor mechanism of action and physiological significance. Prog Neurobiol. 1992;38: 379–394. doi:10.1016/0301-0082(92)90025-A

68. Canavello PR, Cachat JM, Beeson EC, Laffoon AL, Grimes C, Haymore WAM, et al. Measuring Endocrine (Cortisol) Responses of Zebrafish to Stress. In: Kalueff AV, Cachat JM, editors. Zebrafish Neurobehavioral Protocols. Totowa, NJ: Humana Press; 2011. pp. 135–142. doi:10.1007/978-1-60761-953-6_11

69. Bridson D, Forman L. The Herbarium Handbook. U.K.: Kew, Royal Botanic Gardens.: Richmond; 1989.

70. Chang M, Qiu L, Green PS, Wei Z. Oleaceae. In: Wu Z-Y, Raven PH, editors. Flora of China. Missouri Botanical Garden Press; 1996. pp. 272–319.

71. Green PS. Jasminum L. Flora of Thailand. Forest Herbarium, Royal Forest Department; 2000. pp. 308–340.

72. Chalermglin P. Jasmines in Thailand. Amarin Printing and Publishing Public; 2013.

73. Vinatoru M, Mason TJ, Calinescu I. Ultrasonically assisted extraction (UAE) and microwave assisted extraction (MAE) of functional compounds from plant materials. TrAC Trends Anal Chem. 2017;97: 159–178. doi:10.1016/j.trac.2017.09.002

74. Ong ES, Cheong JSH, Goh D. Pressurized hot water extraction of bioactive or marker compounds in botanicals and medicinal plant materials. J Chromatogr A. 2006;1112: 92–102. doi:10.1016/j.chroma.2005.12.052

75. National Institute of Standards and Technology. NIST Tandem mass spectral library. In: NIST 23 Tandem Mass Spectral Libraries [Internet]. 2023 [cited 5 Oct 2023]. Available: https://chemdata.nist.gov/dokuwiki/doku.php?id=chemdata:start

76. Ochocki AJ, Kenney JW. A gelatin-based feed for precise and non-invasive drug delivery to adult zebrafish. J Exp Biol. 2023;226: jeb245186. doi:10.1242/jeb.245186

77. McMillan StephanieC, Géraudie J, Akimenko M-A. Pectoral Fin Breeding Tubercle Clusters: A Method to Determine Zebrafish Sex. Zebrafish. 2015;12: 121–123. doi:10.1089/zeb.2014.1060

78. Mathis A, Mamidanna P, Cury KM, Abe T, Murthy VN, Mathis MW, et al. DeepLabCut: markerless pose estimation of user-defined body parts with deep learning. Nat Neurosci. 2018;21: 1281–1289. doi:10.1038/s41593-018-0209-y

79. Goodwin NL, Choong JJ, Hwang S, Pitts K, Bloom L, Islam A, et al. Simple Behavioral Analysis (SimBA) as a platform for explainable machine learning in behavioral neuroscience. Nat Neurosci. 2024;27: 1411–1424. doi:10.1038/s41593-024-01649-9

80. Blaser R, Gerlai R. Behavioral phenotyping in zebrafish: Comparison of three behavioral quantification methods. Behav Res Methods. 2006;38: 456–469. doi:10.3758/BF03192800

81. Porfiri M, Karakaya M, Sattanapalle RR, Peterson SD. Emergence of in-line swimming patterns in zebrafish pairs. Flow. 2021;1: E7. doi:10.1017/flo.2021.5

82. Parke DV, Rahman KHMQ, Walker R. The Absorption, Distribution and Excretion of Linalool in the Rat. Biochem Soc Trans. 1974;2: 612–615. doi:10.1042/bst0020612

83. Shi F, Zhao Y, Firempong CK, Xu X. Preparation, characterization and pharmacokinetic studies of linalool-loaded nanostructured lipid carriers. Pharm Biol. 2016;54: 2320–2328. doi:10.3109/13880209.2016.1155630

84. R Core Team. R: A Language and Environment for Statistical Computing. In: The R Project for Statistical Computing [Internet]. 2023. Available: https://www.R-project.org/

85. Wickham H. ggplot2: Elegant Graphics for Data Analysis. Springer-Verlag New York; 2016. Available: https://ggplot2.tidyverse.org/

86. Dunnett CW. A Multiple Comparison Procedure for Comparing Several Treatments with a Control. J Am Stat Assoc. 1955;50: 1096–1121. doi:10.1080/01621459.1955.10501294

87. Benjamini Y, Hochberg Y. Controlling the False Discovery Rate: A Practical and Powerful Approach to Multiple Testing. J R Stat Soc Ser B Methodol. 1995;57: 289–300.

88. Cohen J. Statistical Power Analysis for the Behavioral Sciences. 2nd ed. New York: Routledge; 1988. doi:10.4324/9780203771587

