## Supplemental Figure 1 for "The effects of Arabian jasmine on zebrafish behavior depends on strain, sex, and personality"

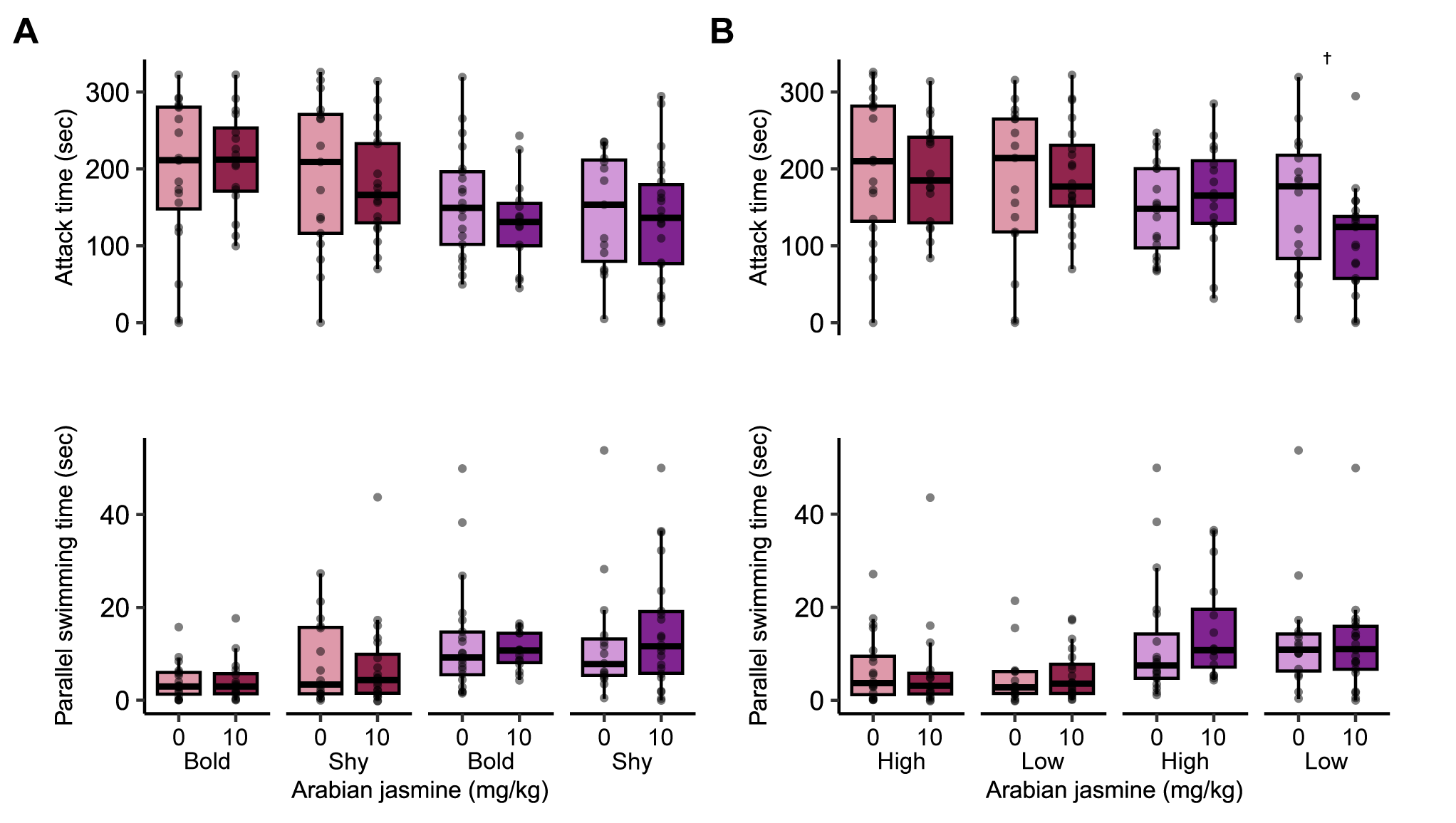

**Figure S1 Influence of Arabian jasmine on aggressive behaviors based on personality.** Social behaviors (attack time and parallel swim time) on day 2 based on (A) boldness and (B) activity in animals given either vehicle or 10 mg kg^-1^ Arabian jasmine. Boxplots indicate median (center line), interquartile range (box ends), and hinge±1.5 times the interquartile range (whiskers). †P<0.10 from pairwise t-test with FDR correction within group. Bold female: *n*=19 vehicle; *n*=16 Arabian jasmine. Shy female: *n*=17 vehicle; *n*=20 Arabian jasmine. Bold male: *n*=21 vehicle; *n*=15 Arabian jasmine. Shy male: *n*=15 vehicle; *n*=22 Arabian jasmine. High-active female: *n*=19 vehicle; *n*=16 Arabian jasmine. Low-active female: *n*=17 vehicle; *n*=20 Arabian jasmine. High-active male: *n*=20 vehicle; *n*=16 Arabian jasmine. Low-active male: *n*=16 vehicle; *n*=21 Arabian jasmine.
